## Supplementary File 1 for "*Corynebacterium glutamicum* regulation beyond transcription: Organizing principles and reconstruction of an extended regulatory network incorporating regulations mediated by small RNA and protein-protein interactions"

**Supplementary figures**

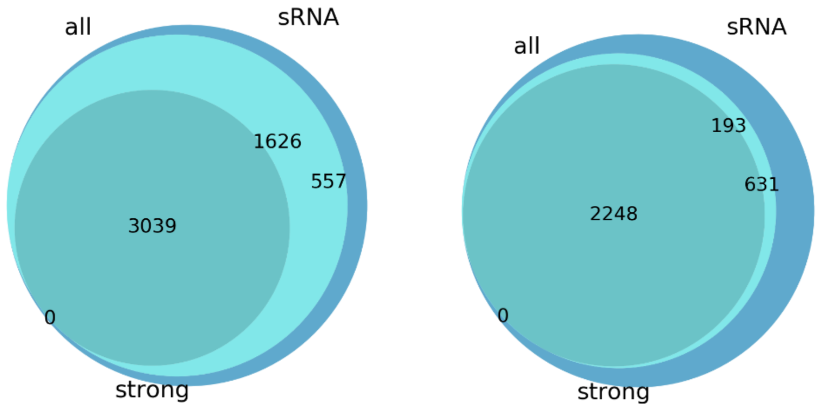


**Supplementary figure 1.** The overlap between the tree network models of *C. glutamicum* for nodes (left) and interactions (right).

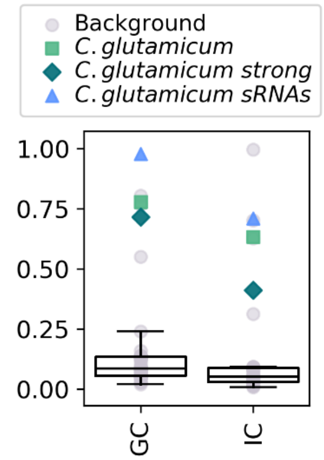


**Supplementary figure 2.** Distribution of the genomic coverage (GC) and interactions coverage (IC) for the non-redundant set of regulatory networks from Abasy Atlas. The data points for the three *C. glutamicum* networks reported in this work are highlighted. Please note that the GC for the sRNA network is inflated by the sRNA nodes (545/3072) although there are still 86 protein-coding genes more than in the *all evidence* network.

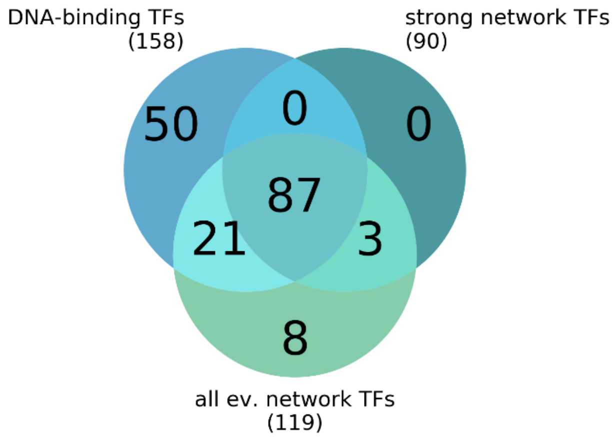


**Supplementary figure 3.** Overlap of transcription factors between the two transcriptional regulatory networks and the complete set of DNA-binding transcription factors in *C. glutamicum*.

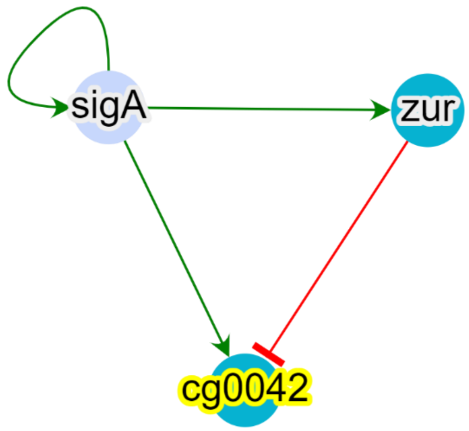


**Supplementary figure 4.** Interaction *zur-cg0042* part of the *C. glutamicum strong* network also recovered from *S. coelicolor*.

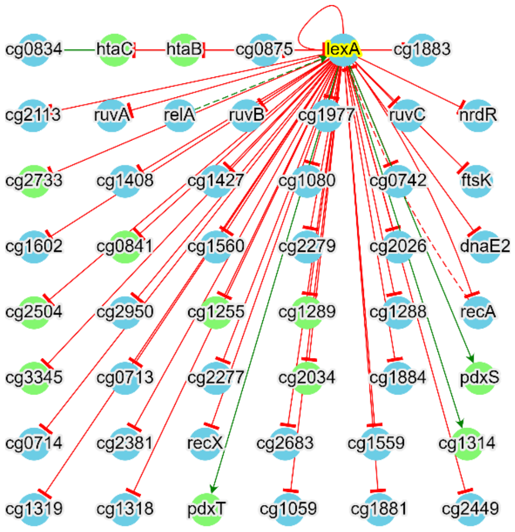


**Supplementary figure 5.** LexA auto-regulation part of the *C. glutamicum* network strongly supported and recovered from *B. subtilis*.

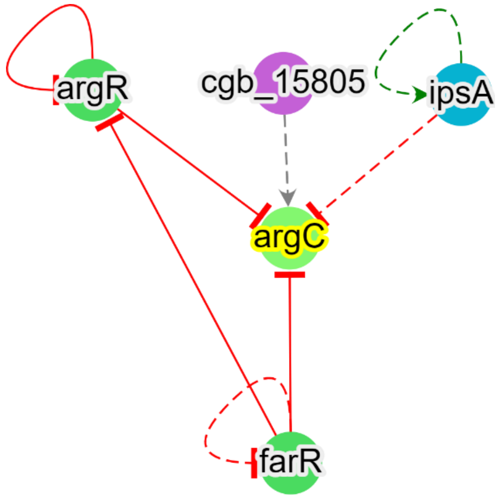


**Supplementary figure 6.** Interaction *argR*-*argC* part of the *C. glutamicum* strongly supported and recovered from *E. coli*.

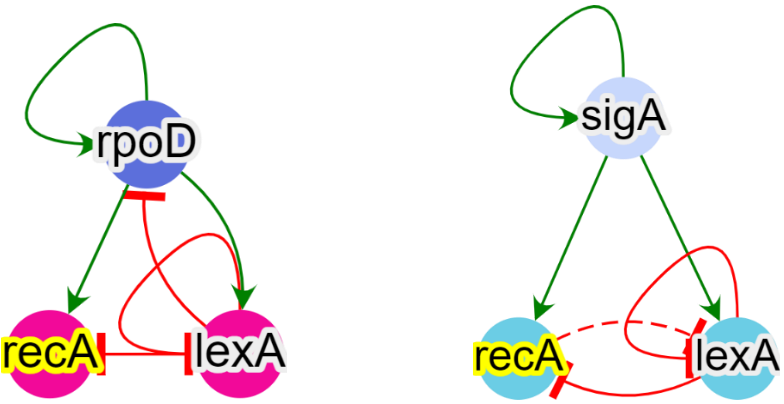


**Supplementary figure 7.** The interaction *lexA-recA* was recovered from *E. coli* (left) and is already part of the *C. glutamicum* network (right) strongly supported.

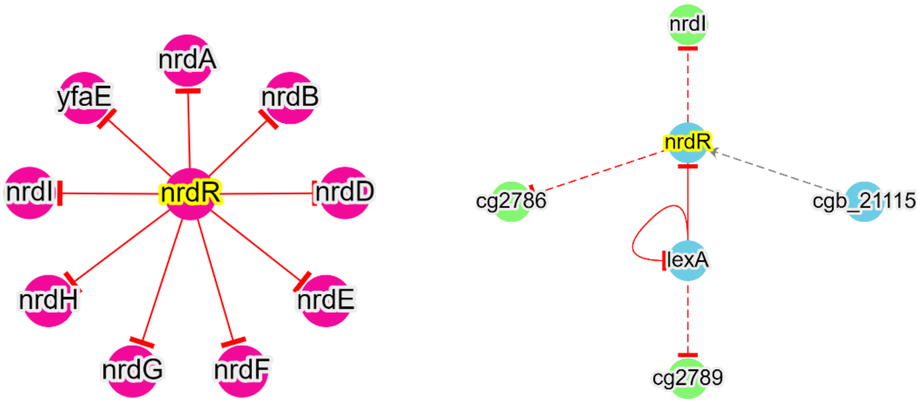


**Supplementary figure 8.** The interaction *nrdR-nrdI* was recovered from *E. coli* (left) and is already part of the *C. glutamicum* network (right) with non-strong evidence supporting it.

**Supplementary tables**

**Supplementary Table 1.** Interactions from other *C. glutamicum* strains or not mapping to a cg number. Please consider this is not an exhaustive list of regulations for other strains.

| TF | TG | Effect |
|---|---|---|
| *cg3247* | *NCgl2845* | + |
| *cg3247* | *NCgl1729* | + |
| *cg0702* | *cg1652* | - |
| *cg0702* | *cg1583* | - |
| *cg0702* | *cg3378* | - |
| *cg0702* | *cg0400* | + |
| *cg0702* | *NCgl0166* | + |
| *cg0702* | *cg1582* | - |
| *cg0702* | *cg3387* | + |
| *cg0702* | *NCgl0484* | + |
| *cg0702* | *cg0771* | + |
| *cg0702* | *cg3327* | - |
| *cg3224* | *NCgl1861* | - |
| *cg0702* | *cg0344* | + |
| *cg3247* | *NCgl2113* | + |
| *cg0702* | *NCgl2942* | + |
| *cg0702* | *NCgl2893* | + |
| *cg0702* | *cg1215* | - |
| *cg0702* | *cg3375* | + |
| *cg0702* | *NCgl0746* | - |
| *cg0146* | *NCgl18* | + |
| *cg0702* | *cg1528* | - |
| *cg0702* | *NCgl0608* | + |
| *cg0702* | *cg2252* | + |
| *cg3224* | *NCgl2817* | - |
| *cg0702* | *cg0345* | + |
| *cg0702* | *cg1581* | - |
| *cg2831* | *NCgl1806* | + |
| *cg3224* | *NCgl0361* | + |
| *cg0702* | *cg1584* | - |
| *cg3247* | *NCgl2738* | - |
| *cg3247* | *NCgl2237* | - |
| *cg0702* | *NCgl0580* | + |
| *cg3247* | *NCgl2861* | + |
| *cg0702* | *NCgl2970* | + |
| *cg0702* | *cg0661* | - |
| *cg3247* | *NCgl1758* | + |

| | | |
|---|---|---|
| *cg0702* | *cg0018* | + |
| *cg3247* | *NCgl0404* | + |
| *cg0702* | *cg1214* | - |
| *cg3224* | *NCgl0360* | + |
| *cg3247* | *NCgl2858a* | + |
| *cg3247* | *NCgl1780* | + |
| *cg0702* | *cg1120* | + |
| *cg0702* | *cg3145* | + |
| *cg0702* | *cg0197* | + |
| *cg0702* | *cg0318* | + |
| *cg2831* | *NCgl1812* | - |
| *cg2831* | *NCgl1783* | + |
| *cg3247* | *NCgl1750* | + |
| *cg0702* | *NCgl0638* | + |

**Supplementary Table 2**. Interactions recovered from *S. coelicolor, B. subtilis,* and *E. coli*. The *Average rank* is the averaged ranking position between the three inferences with the three motif finding tools (smaller is better). The *Status* can take three values: 1) *network* for interactions that have been already experimentally validated and are part of one of the *C. glutamicum* networks; 2) *potential new TG* for the interactions mediated by a TF that is already a TF in one of the networks, but its regulation of the TG expression has not been experimentally validated; and 3) *new TF* for interactions mediated by TFs with uncharacterized regulons, hence those that are not included in the current networks.

| TF | TG | Average rank | Status |
|---|---|---|---|
| Recovered from *S. coelicolor* | | | |
| *cg0484* | *cg2261* | 10.0 | new TF |
| *cg1585* | *cg0850* | 42.333 | potential new TG |
| *cg2502* | *cg0042* | 50.0 | network |
| *cg1585* | *cg2305* | 51.0 | potential new TG |
| *cg3202* | *cg2117* | 54.333 | potential new TG |
| *cg0484* | *cg2846* | 57.333 | new TF |
| *cg2502* | *cg0991* | 59.333 | potential new TG |
| *cg0484* | *cg1809* | 61.333 | new TF |
| *cg3202* | *cg2929* | 62.0 | potential new TG |
| *cg1585* | *cg0113* | 63.0 | potential new TG |
| *cg0484* | *cg2260* | 65.333 | new TF |
| *cg1486* | *cg1487* | 67.0 | network |
| *cg0484* | *cg2485* | 67.333 | new TF |
| *cg1585* | *cg1580* | 68.0 | network |
| *cg2109* | *cg2109* | 69.0 | network |

| | | | |
|---|---|---|---|
| *cg1585* | *cg0303* | 71.0 | potential new TG |
| *cg1486* | *cg2383* | 71.0 | potential new TG |
| *cg1585* | *cg1586* | 72.0 | potential new TG |
| *cg1585* | *cg2261* | 73.333 | potential new TG |
| *cg1098* | *cg1098* | 74.333 | new TF |
| *cg0484* | *cg2513* | 76.667 | new TF |
| *cg0313* | *cg0313* | 77.0 | network |
| *cg1486* | *cg1486* | 83.667 | potential new TG |
| *cg3202* | *cg3202* | 85.667 | network |
| Recovered from *B. subtilis* | | | |
| *cg2114* | *cg2114* | 3.333 | network |
| *cg2114* | *cg2141* | 11.667 | network |
| *cg2114* | *cg1996* | 15.333 | potential new TG |
| *cg2624* | *cg2803* | 25.0 | potential new TG |
| *cg1585* | *cg1580* | 27.333 | network |
| *cg2624* | *cg1142* | 29.667 | potential new TG |
| *cg1585* | *cg1582* | 39.333 | network |
| *cg2516* | *cg3100* | 39.667 | potential new TG |
| *cg3097* | *cg0113* | 40.0 | potential new TG |
| *cg1817* | *cg1814* | 40.667 | potential new TG |
| *cg1817* | *cg1815* | 42.333 | network |
| *cg2516* | *cg2514* | 44.667 | potential new TG |
| *cg2114* | *cg1401* | 47.333 | potential new TG |
| *cg1817* | *cg1817* | 48.0 | network |
| *cg2516* | *cg3099* | 49.667 | potential new TG |
| *cg2114* | *cg1560* | 49.667 | network |
| *cg2114* | *cg0976* | 50.667 | potential new TG |
| Recovered from *E. coli* | | | |
| *cg1585* | *cg1580* | 19.0 | network |
| *cg2114* | *cg2114* | 62.667 | network |
| *cg2114* | *cg2141* | 64.0 | network |
| *cg0350* | *cg2166* | 115.333 | potential new TG |
| *cg0350* | *cg3395* | 164.333 | potential new TG |
| *cg2899* | *cg2637* | 169.0 | new TF |
| *cg3224* | *cg2559* | 190.667 | potential new TG |
| *cg0350* | *cg1568* | 195.0 | network |
| *cg1585* | *cg3004* | 201.0 | potential new TG |
| *cg0350* | *cg2429* | 209.667 | network |
| *cg0350* | *cg1257* | 211.333 | potential new TG |
| *cg0350* | *cg0229* | 212.0 | network |
| *cg0001* | *cg0001* | 215.667 | new TF |
| *cg1327* | *cg1327* | 217.0 | new TF |

| | | | |
|---|---|---|---|
| *cg2502* | *cg0591* | 224.0 | potential new TG |
| *cg0350* | *cg2175* | 229.667 | potential new TG |
| *cg0350* | *cg2126* | 232.0 | potential new TG |
| *cg1585* | *cg2167* | 235.0 | potential new TG |
| *cg0350* | *cg3068* | 243.667 | potential new TG |
| *cg0350* | *cg2870* | 243.667 | potential new TG |
| *cg0350* | *cg1145* | 244.0 | network |
| *cg1327* | *cg3141* | 250.667 | new TF |
| *cg2502* | *cg2782* | 253.667 | potential new TG |
| *cg2936* | *cg2933* | 254.333 | network |
| *cg2502* | *cg3237* | 255.0 | potential new TG |
| *cg0350* | *cg3308* | 256.0 | potential new TG |
| *cg0350* | *cg2102* | 263.0 | potential new TG |
| *cg0350* | *cg1492* | 266.667 | potential new TG |
| *cg0001* | *cg1525* | 273.667 | new TF |
| *cg2112* | *cg2787* | 281.667 | network |
| *cg1425* | *cg0001* | 283.333 | potential new TG |
| *cg2502* | *cg2183* | 285.0 | potential new TG |
| *cg0350* | *cg2841* | 286.0 | potential new TG |
| *cg1585* | *cg2166* | 287.333 | potential new TG |
| *cg0350* | *cg1790* | 290.333 | network |
| *cg0350* | *cg3340* | 290.667 | potential new TG |
| *cg2502* | *cg2502* | 291.0 | potential new TG |
| *cg0350* | *cg3423* | 291.0 | potential new TG |
| *cg0350* | *cg1586* | 293.0 | potential new TG |
| *cg1327* | *cg1656* | 293.667 | new TF |
| *cg1585* | *cg2178* | 298.0 | potential new TG |
| *cg1585* | *cg0229* | 299.0 | network |
| *cg0350* | *cg0350* | 299.0 | network |
| *cg0350* | *cg0067* | 300.667 | potential new TG |
| *cg2114* | *cg2489* | 303.0 | potential new TG |
| *cg0350* | *cg0953* | 304.333 | potential new TG |
| *cg1425* | *cg0004* | 309.0 | potential new TG |
| *cg1585* | *cg1588* | 310.0 | potential new TG |
| *cg2112* | *cg2781* | 313.0 | potential new TG |
| *cg1585* | *cg1586* | 314.333 | potential new TG |
| *cg1585* | *cg1814* | 317.333 | network |
| *cg1327* | *cg1355* | 320.0 | new TF |
| *cg2112* | *cg2789* | 327.667 | network |
| *cg0001* | *cg0004* | 328.0 | new TF |
| *cg1327* | *cg2856* | 329.0 | new TF |
| *cg1425* | *cg0306* | 331.0 | potential new TG |

| | | | |
|---|---|---|---|
| *cg2899* | *cg2899* | 331.667 | new TF |
| *cg0350* | *cg0699* | 333.667 | potential new TG |
| *cg1425* | *cg0005* | 334.667 | potential new TG |
| *cg0350* | *cg0673* | 340.667 | potential new TG |
| *cg0350* | *cg3336* | 345.333 | potential new TG |
| *cg0350* | *cg1163* | 351.667 | potential new TG |
| *cg1327* | *cg1383* | 362.333 | new TF |
| *cg0350* | *cg2932* | 362.667 | potential new TG |
| *cg0001* | *cg1550* | 370.333 | new TF |
| *cg0001* | *cg0005* | 374.333 | new TF |
| *cg2114* | *cg1602* | 375.0 | network |
| *cg1327* | *cg2891* | 376.667 | new TF |
| *cg0350* | *cg2178* | 379.0 | potential new TG |
| *cg0350* | *cg3096* | 379.667 | network |
